# Protein folding success depends on the direction and speed of polypeptide chain appearance

**DOI:** 10.1101/2025.06.30.662406

**Authors:** Iker F. Soto-Santarriaga, Patricia L. Clark

## Abstract

The failure of proteins to fold correctly to their native, functional structures is the molecular basis of many human diseases. Every protein is introduced to the cellular environment from N-to C-terminus as it is synthesized, which can enable the N-terminus to begin folding before the C-terminus appears. Here we show that direction and speed of appearance alone can significantly affect whether a protein will fold to its native structure, versus misfold. This sensitivity demonstrates that folding energy landscapes are sufficiently rugged that conformations adopted by the first portion of the protein to appear can significantly impact subsequent folding steps. Despite high structural similarity, homologous proteins had different sensitivities to vectorial appearance, indicating the native topology is insufficient to determine contours of the energy landscape.

---

As peptide bonds are formed by the ribosome, the growing N-terminus of every newly synthesized protein first passes through the narrow (1-2 nm diameter) ribosome exit tunnel^1,2^ before emerging into the cytoplasm, enabling the N-terminal portion of the protein to begin folding prior to the appearance of the C-terminus^3–5^. Similarly, during or shortly after protein synthesis many proteins are translocated from one end to the other across a lipid membrane into another subcellular compartment, where folding can also commence vectorially^6,7^. Yet despite the ubiquity of vectorial appearance and its potential influence on protein folding mechanisms, most of our current understanding of protein folding pathways^8–10^, including investigations of the effects of disease-causing mutations^11,12^, is instead derived from diluting full-length proteins out of a chemical denaturant. Dilution enables folding to start with interactions formed between any two parts of the protein, rather than providing a temporal advantage to one end of the protein.

There is some evidence that initiating folding during vectorial appearance can significantly alter the folding outcome. Vectorial folding during protein synthesis can lead to the population of distinct intermediate conformations not populated during *in vitro* refolding, which in turn has been implicated to affect whether a protein will fold correctly or instead misfold^13–16^. Moreover, a direct prediction of the importance of folding during vectorial appearance is that modulating the rate of translation elongation by the ribosome should affect protein folding outcomes, which has now been reported for several proteins^17–23^. More broadly, only a small subset of proteins in the proteome have been shown to fold robustly to their native structure upon dilution from denaturant ^24^. Instead, most misfold and often aggregate^25^. Extensive investigations of misfolding and aggregation mechanisms have revealed that there is typically a large energy barrier between the native and misfolded states^26,27^, highlighting the crucial importance of early conformations to the partitioning between the folding and misfolding minima on a protein folding energy landscape (**Fig. 1a**) ^27–29^.

**Fig. 1.**
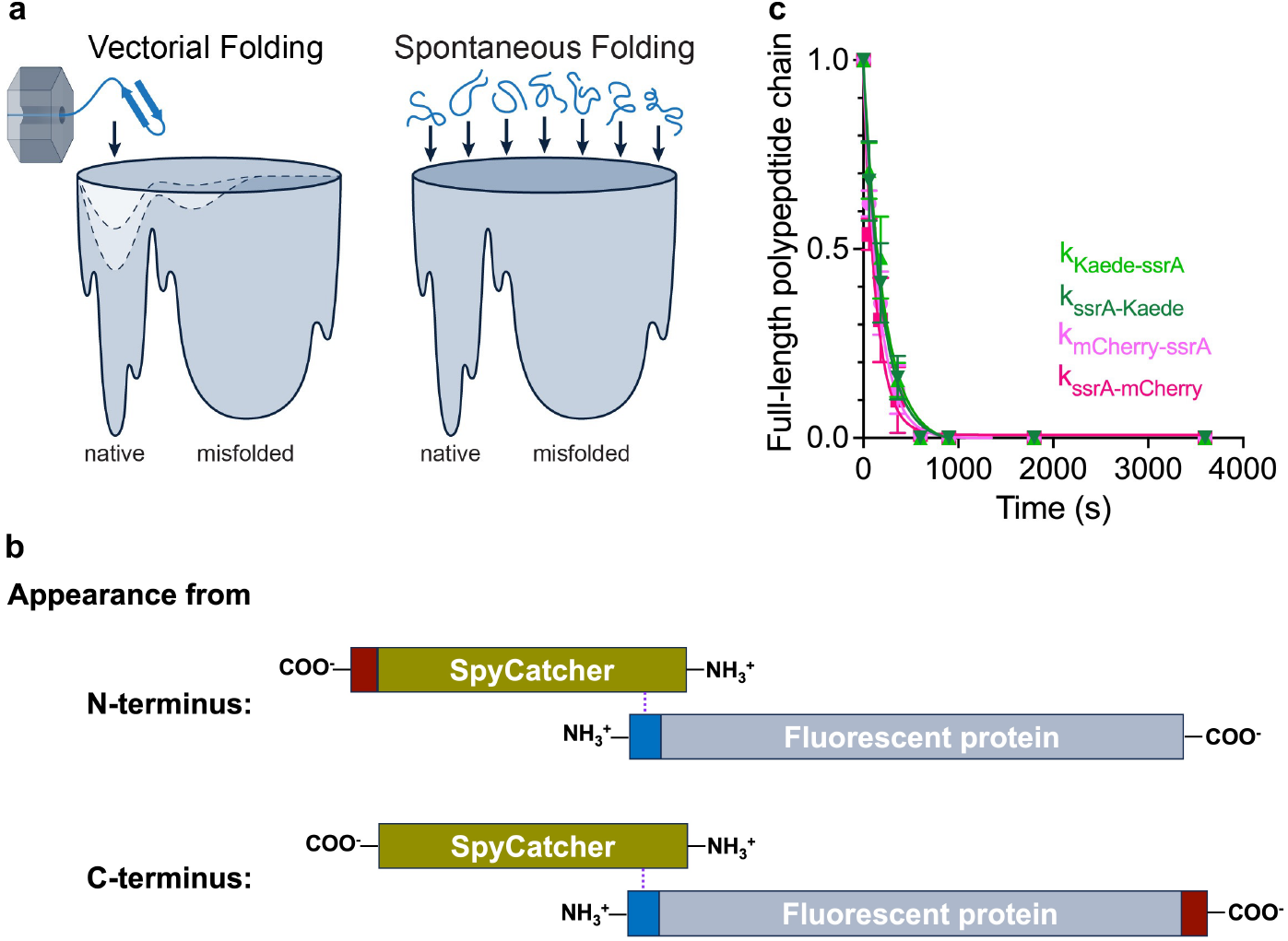
Hypotheses tested and development of ClpX as a tool for initiating protein folding during vectorial appearance. (**a**) Folding during vectorial appearance (*left*) initially restricts access to portions of the energy landscape that require the other terminus of the protein chain (*dashed lines*). This phenomenon could enhance (or suppress; not shown) protein folding efficiency versus dilution of a full-length protein from denaturant (*right*). Note that the energy landscape for the full-length protein remains unchanged regardless of how folding is initiated. (**b**) Design of substrates used to initiate protein folding upon vectorial appearance from the central pore of ClpX. SpyCatcher (*yellow*) was covalently attached to SpyTag (blue) at the N-terminus of each fluorescent protein (FP; *grey*) via an isopeptide bond (*dashed line*). The direction of appearance of each FP from ClpX was specified by encoding the ssrA tag (*red*) at either the C-terminus of SpyCatcher (to initiate appearance from N-terminus; *top*) or the C-terminus of the FP (to initiate appearance from C-terminus; *bottom*). (**c**) The degradation rate of substrate proteins diluted from chemical denaturant is indistinguishable regardless of the FP sequence, tag location or direction of vectorial appearance; mCherry-ssrA (light pink), ssrA-mCherry (dark pink), Kaede-ssrA (light green), ssrA-Kaede (dark green). Degradation rate was measured by quantifying SDS-PAGE band intensity; see Methods. Error bars represent the standard error of the mean (SEM) for three independent experiments.

However, an ongoing challenge to understanding the impact of vectorial appearance on protein folding mechanisms is the complexity of the cellular environment. For example, the cytoplasm also includes molecular chaperones, which themselves can contribute to the successful folding of some proteins^30–32^. Likewise, although synonymous codon substitutions were long thought to be phenotypically “silent” and are often used to manipulate the rate of translation elongation, these substitutions have now been shown to affect many mechanisms related to gene expression, beyond nascent protein folding^33,34^. An additional challenge is that even a minimal *in vitro* translation system must include >50 different proteins and >20 RNA molecules, creating a complex background that complicates measurements of nascent protein folding. Finally, a rigorous analysis of the impact of vectorial appearance would include a comparison of folding outcomes when vectorial appearance is initiated from the C-versus N-terminus, an important control that is incompatible with protein synthesis, as ribosomes synthesize proteins only from N-to-C-terminus. The asymmetry of the peptide bond means that C-to-N-terminal appearance cannot be replicated simply by placing an amino acid sequence in the opposite order, as disrupting the relationship between peptide bond direction and amino acid sequence can radically reshape the energy landscape for folding^35^. Similarly, circular permutation, which changes the order of appearance of protein segments by covalently connecting the natural N- and C-terminus and introducing new termini elsewhere^36,37^ can also significantly perturb the energy landscape for folding^38,39^.

## Tunable, unidirectional vectorial appearance through the central pore of ClpX

To isolate and evaluate the impact of vectorial appearance on protein folding mechanisms, we exploited the *E. coli* AAA+ protein ClpX, which uses the energy of ATP hydrolysis to mechanically unfold and translocate proteins through its narrow central pore^40,41^. The diameter of the ClpX pore is similar to the diameter of the ribosome exit tunnel^42,43^, and the ClpX translocation rate is similar to the elongation rate of the *E. coli* ribosome^44–46^. Although translocation through ClpX is unidirectional, the direction of translocation can be specified by attaching a ClpX recognition sequence to either the N- or C-terminus of the substrate protein^47^.

As substrates to test the impact of vectorial appearance on folding outcome, we selected two homologous β-barrel fluorescent proteins (FPs), mCherry and Kaede^48,49^. It was previously shown that, for GFP, co-translational folding intermediates fold to native structure much more efficiently than upon dilution from denaturant, which tends to lead to misfolding^14,50^. After FP chromophore formation, which is irreversible, fluorescence emission occurs only when the FP adopts its native structure^51,52^. Consistent with previous reports that FPs are prone to misfolding upon dilution from a chemical denaturant^14,50,53^, we observed low yields for the refolding of either mCherry or Kaede upon dilution from guanidine hydrochloride (GdmCl), measured as low recovery of fluorescence emission (**Fig. S1**).

We first used a ClpXP degradation assay^54^ to test whether FPs tagged with the well-characterized ClpX-recognition ssrA motif (X_7_-ALAA)^55^ are readily translocated through the central pore of ClpX from either the N- or C-terminus. ClpP is the partner protease of ClpX, and degradation of ssrA-tagged substrate proteins by ClpP requires ATP-dependent translocation of substrate proteins through the central pore of ClpX^40^. For C-to-N-terminal translocation, we genetically encoded an ssrA tag at the C-terminus of each FP (**Fig. 1b**). N-terminal tagging required attaching the ssrA sequence “backwards” to the protein N-terminus, to retain the free carboxyl group and C-terminal alanine residues essential for ClpX recognition^55^. Previously, maleimide coupling was used for N-terminal ssrA attachment^47^, however this restricts the selection of substrate proteins to those that lack cysteine residues. We instead genetically encoded SpyTag at the N-terminus of each FP, creating a recognition sequence for the small protein SpyCatcher (SC), which forms an isopeptide bond between Lys_31_ of SpyCatcher and Asp_8_ of SpyTag^56^. We then genetically encoded the ssrA tag at the C-terminus of SpyCatcher, purified it, and incubated it with each SpyTagged FP. This approach enabled the purification of each FP with essentially 100% attachment of the ssrA tag at the N-terminus (**Fig. 1b & S2**). Prior to translocation through ClpX, each tagged FP was denatured in GdmCl, to avoid the possibility of residual native structure slipping through the central pore of ClpX, as such structure could potentially bias the folding outcome. Chemical denaturation prior to dilution into the presence of ClpXP and ATP led to complete degradation of both FPs at the same rate when tagged with ssrA at either the N- or C-terminus (**Fig. 1c**). Degradation was dependent on the presence of ATP, ClpX and the ssrA tag, consistent with tagged substrate protein translocation through the central pore of ClpX^54^.

### Vectorial appearance significantly impacts folding yield

When the ClpP protease was omitted, vectorial appearance from ClpX led to a time-dependent recovery of FP fluorescence emission, instead of degradation, indicating that FPs emerging vectorially from the distal side of ClpX are capable of refolding (**Fig. 2**). Our expectation was that, in this multiple-turnover experiment, the fluorescence intensity plateau represents the establishment of a steady-state between refolding and subsequent rounds of ClpX-mediated unfolding. Consistent with this interpretation, addition of a competitive inhibitor (a 60-fold excess of an alternative ssrA-tagged substrate protein) led to a further enhancement in refolding yield (**Fig. S3**).

**Figure 2.**
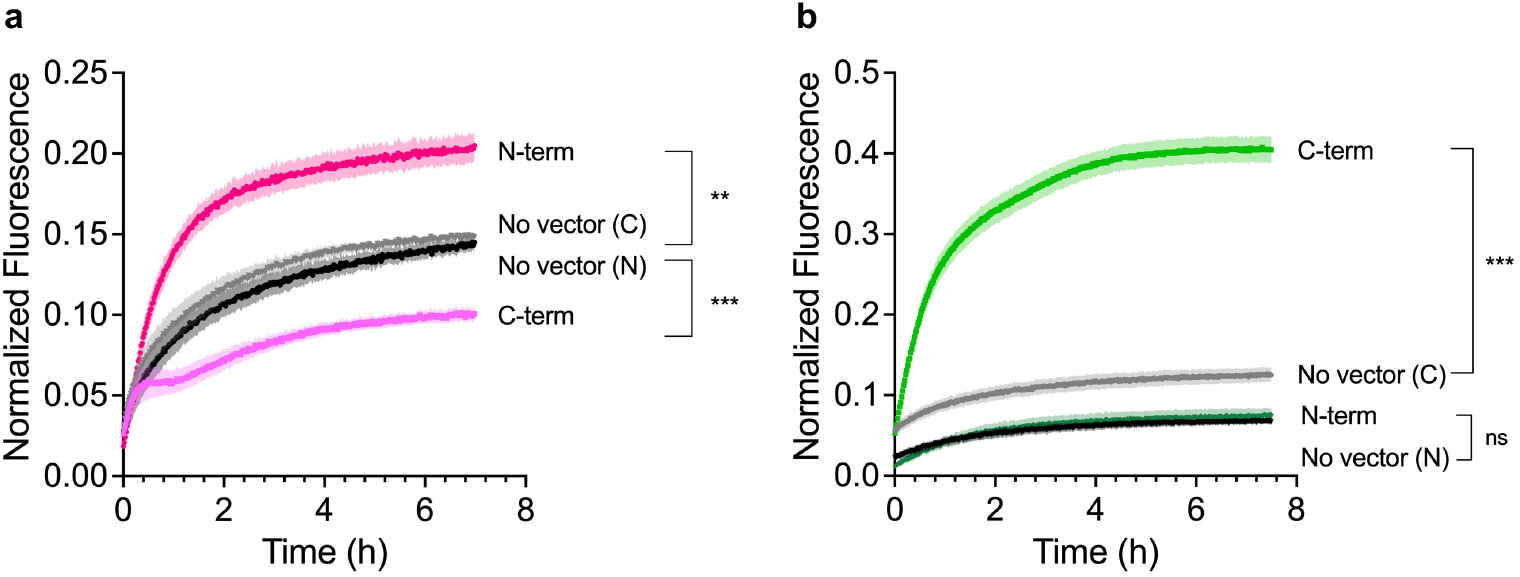
Direction of protein appearance affects folding outcome. (**a**) N-terminal appearance of mCherry (*dark pink*) significantly enhanced folding, whereas C-terminal appearance (*light pink*) significantly suppressed folding compared to the no vector condition (*black* and *grey*). (**b**) C-terminal appearance of Kaede (*light green*) significantly enhanced folding compared to no vector (*grey*), whereas appearance from the N-terminus (*dark green*) was indistinguishable from no vector (*black*). Note that the placement of the ssrA tag alone led to modest differences in Kaede folding even in the absence of vectorial appearance. For all experiments, points represent the mean of four independent experiments and shading correspond to SEM. Fluorescence emission intensity was normalized to the signal for an equivalent concentration of native (never unfolded) protein. Differences between the fluorescence intensities at the final time point of each curve were analyzed using the student’s t-test (**, p<0.01; ***, p<0.001).

Folding outcomes upon vectorial appearance from ClpX were significantly different from dilution from denaturant alone and depended on the direction of vectorial appearance. For mCherry, appearance from the N-terminus significantly increased the folding yield versus dilution from denaturant alone, whereas appearance from the C-terminus significantly suppressed folding to the native state (**Fig. 2a**). These results indicate that vectorial appearance leads to the formation of distinct folding intermediates depending on which terminus appeared first, with those formed upon vectorial appearance from the N-terminus being more likely to support formation of the native structure than the folding intermediates formed when the C-terminus appears first. Despite sharing 50% sequence identity with mCherry, Kaede exhibited distinctly different sensitivity to vectorial appearance. Appearance from the C-terminus led to a nearly four-fold increase in Kaede folding yield, whereas N-terminal appearance had no impact versus dilution from denaturant (**Fig. 2b**). These results indicate that although the sequences of mCherry and Kaede are sufficiently similar as to yield highly similar native structures, and both proteins have folding mechanisms that are sensitive to vectorial appearance, their sequence disparities lead to distinctly different energy landscapes for folding. As an initial exploration of this marked difference in vectorial folding behavior, we analyzed where the greatest sequence differences occur in an alignment of mCherry and Kaede (**Fig. S4a**). The greatest sequence disparity is in the C-terminal 20% of the alignment (**Fig. S4b**), consistent with the marked difference in folding behavior upon C-to-N-terminal appearance.

The results described above are consistent with studies of small proteins that refold reversibly, which revealed that the refolding rates of homologs can differ by several orders of magnitude^57^ and populate distinct intermediates^58^. Similarly, we observed significant differences in FP folding kinetics upon vectorial appearance. Specifically, the enhancement in Kaede folding yield upon appearance from the C-terminus was correlated with the appearance of a second, faster exponential change in the observable time frame (*k*_fast_; 7.3×10^−4^ sec^-1^) (**Fig. 2b & Table S1**). A similar faster phase was also observed during N-to-C-terminal appearance of mCherry but was not detected during the low-yield folding (e.g., folding of mCherry upon vectorial appearance from C-to-N-terminus, Kaede folding during N-to-C-terminal appearance, or in the absence of vectorial appearance) (**Fig. 2a & Table S1**).

### Rate of appearance also affects folding yield

Previous work has shown that the rate of translation elongation can be modulated by substitutions between rare and common synonymous codons^59^, and these modulations can alter co-translational folding outcomes^21,22^. To determine if the rate of appearance from ClpX can similarly affect the folding for mCherry or Kaede, we reduced the ClpX translocation rate by using a 1:1 mixture of ATP and ATPγS (**Fig. S5**)^54,60^. In general, slower translocation through ClpX enhanced FP folding yield (**Fig. 3**), and in no case did slowing the rate of appearance reduce folding yield. These results are consistent with a model where slower appearance is generally beneficial, providing additional time for the first part of the protein that appears to adopt a conformation consistent with subsequent rapid and efficient formation of the native structure (**Fig. 4**), before introducing additional portions of the sequence that might instead stabilize off-pathway, misfolded conformations^61^.

**Fig. 3.**
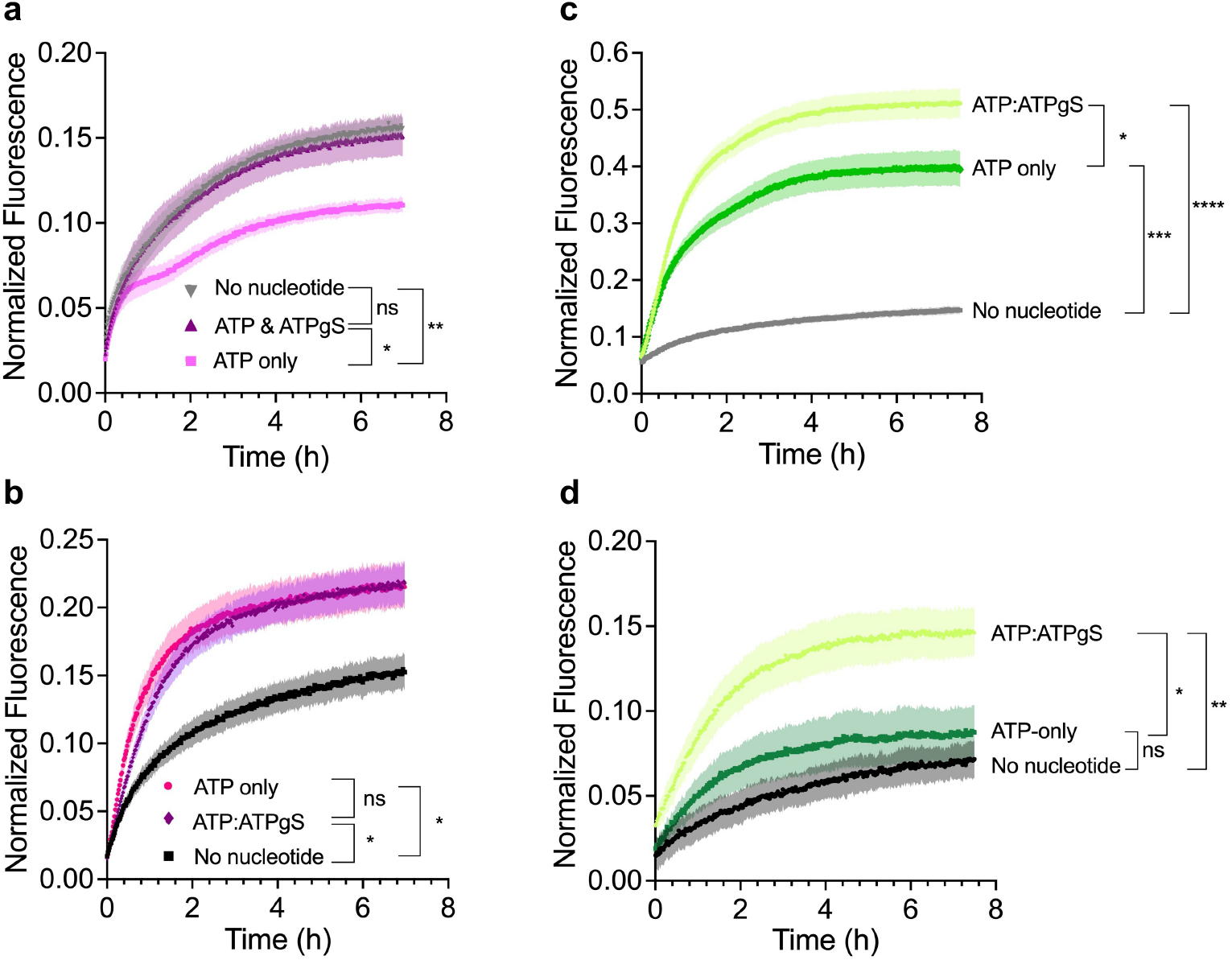
Protein folding outcomes are sensitive to the rate of vectorial appearance. (**a**) Slowing the C-terminal emergence of mCherry (*purple*) reduces the folding suppression seen under the ATP-only condition (*light pink*). (**b**) Slowing the N-terminal emergence of mCherry (*purple*) does not enhance folding beyond the ATP-only condition (*dark pink*). (**c**) Slowing the C-terminal emergence of Kaede (*chartreuse*) improves folding yield over the ATP-only condition (*lime green*). (**d**) Slowing the N-terminal emergence of Kaede increases folding yield beyond the ATP-only condition, though not to the same extent as seen for Kaede-ssrA. For all experiments, points represent the mean of four independent experiments and shading correspond to SEM. Fluorescence emission intensity was normalized to the signal for an equivalent concentration of native (never unfolded) protein. Differences between the fluorescence intensities at the final time point of each curve were analyzed using the student’s t-test (*, p<0.05; **p<0.01; **, p<0.01; ***, p<0.001).

**Fig 4.**
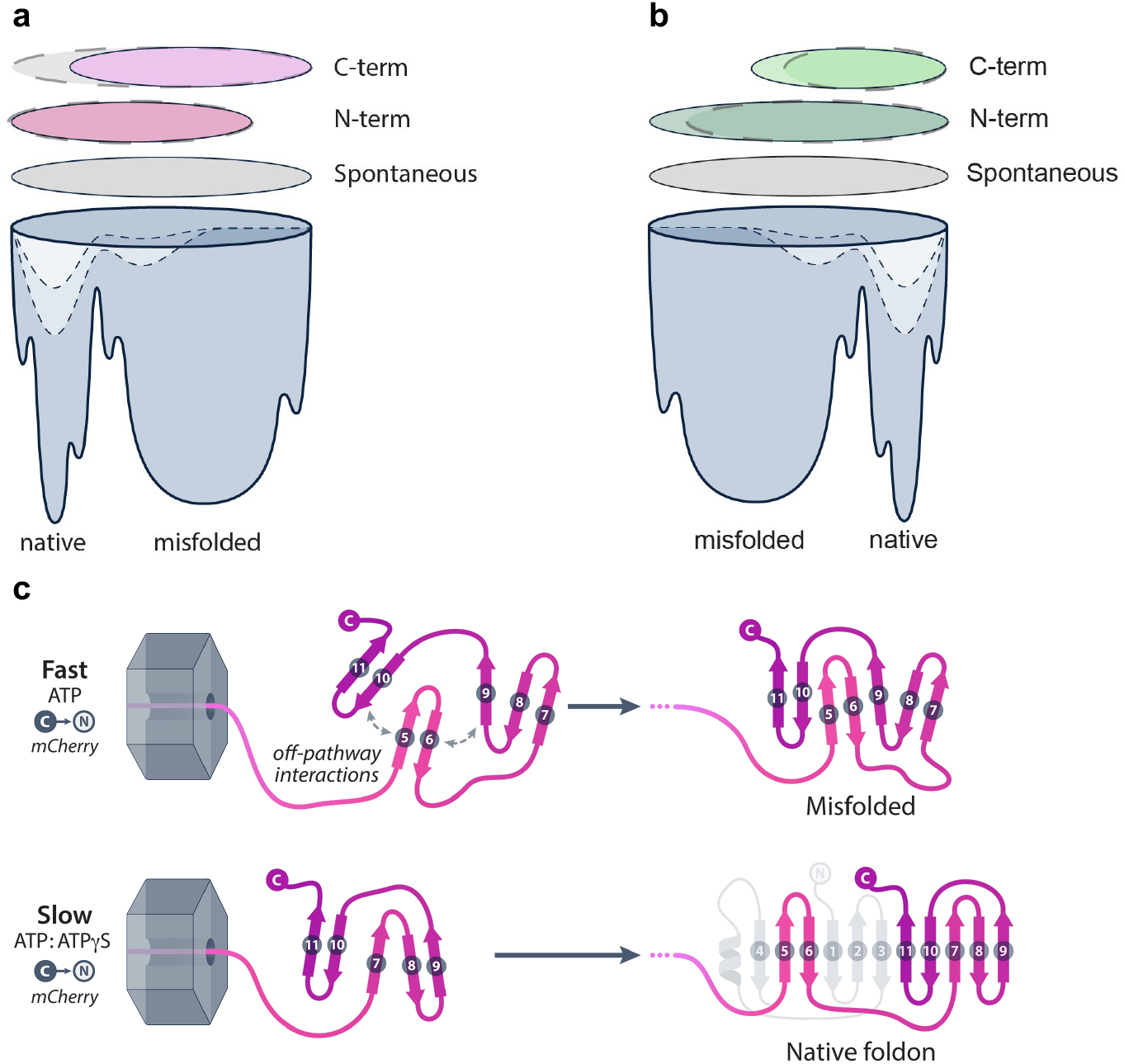
Effects of direction and rate of vectorial appearance on FP folding outcomes. (**a**-**b**) Unlike spontaneous refolding (*gray circles*), where the unfolded protein can immediately access its full conformational landscape, vectorial folding restricts the emerging polypeptide chain of (**a**) mCherry and (**b**) Kaede to specific regions within their conformational space (*pink and green circles, respectively*). This gradual exposure biases the folding pathway and alters the likelihood of reaching the native state versus misfolding. Slowing the rate of vectorial appearance further modulates this conformational bias (*dashed lines*), often enhancing folding efficiency. (**c**) Schematic illustrating how translocation rate may influence folding outcomes. Rapid appearance of mCherry from the C-terminus (*top*) can facilitate non-native contacts that lead to misfolding. In contrast, slowing the emergence of the C-terminus (*bottom*) provides more time for the C-terminus to fold before the appearance of the more N-terminal portions of the protein that can lead to misfolding.

### Residual native structure retained during ClpX translocation can bias folding outcome

The results above demonstrate that FP folding rate and yield is sensitive to the direction and rate of appearance of the protein chain. Yet in all cases, the yield of native protein remained <60%, suggesting that refolding from a fully denatured ensemble can still be inefficient, even when improved by vectorial appearance. Alternatively, ClpX is capable of mechanically unfolding native proteins via its ATP hydrolysis-driven power stroke, prior to translocation ^44,46^. Although the central pore of ClpX is too narrow to accommodate translocation of a native FP^42,43^, small, stable structures, including knots and covalently linked polypeptide chains, have been shown to pass through the central pore of ClpX^62,63^. Similarly, local folding of the nascent protein can start far within the ribosome exit tunnel^64,65^, although it is currently unclear whether these local structures are sufficiently stable to bias subsequent folding steps, after the nascent chain emerges from the confinement of the exit tunnel^66^.

To test whether local native conformational biases can persist during translocation through ClpX, and the extent to which these biases impact the subsequent folding of substrate proteins to their native structures, we assessed FP refolding after ClpX-mediated translocation of native – rather than pre-denatured – ssrA-tagged FPs. The final folding yield was significantly higher for both mCherry and Kaede when pre-denaturation was omitted (**Fig. 5**). Translocation of native FP substrates led to an initial loss of fluorescence that reached a plateau. As above, consistent with this plateau representing a steady-state between ClpX-mediated unfolding and post-translocation refolding, addition of a competitive inhibitor enhanced FP refolding level still further (**Fig. S3**). The discrepancy in folding outcomes for substrate proteins that were pre-denatured versus those that were not is consistent with a model where ClpX-mediated unfolding of native FPs leads to the retention of some local, native-like conformational biases that are sufficiently stable as to enhance the folding yield after vectorial appearance, retaining proteins near the native basin of the folding energy landscape (**Fig. 4**). In contrast, pre-denaturation in GdmCl results in a starting ensemble of FP conformations that is more representative of a random ensemble of all possible starting conformations, including those that are prone to misfolding, which can be suppressed by folding during vectorial appearance.

**Figure 5.**
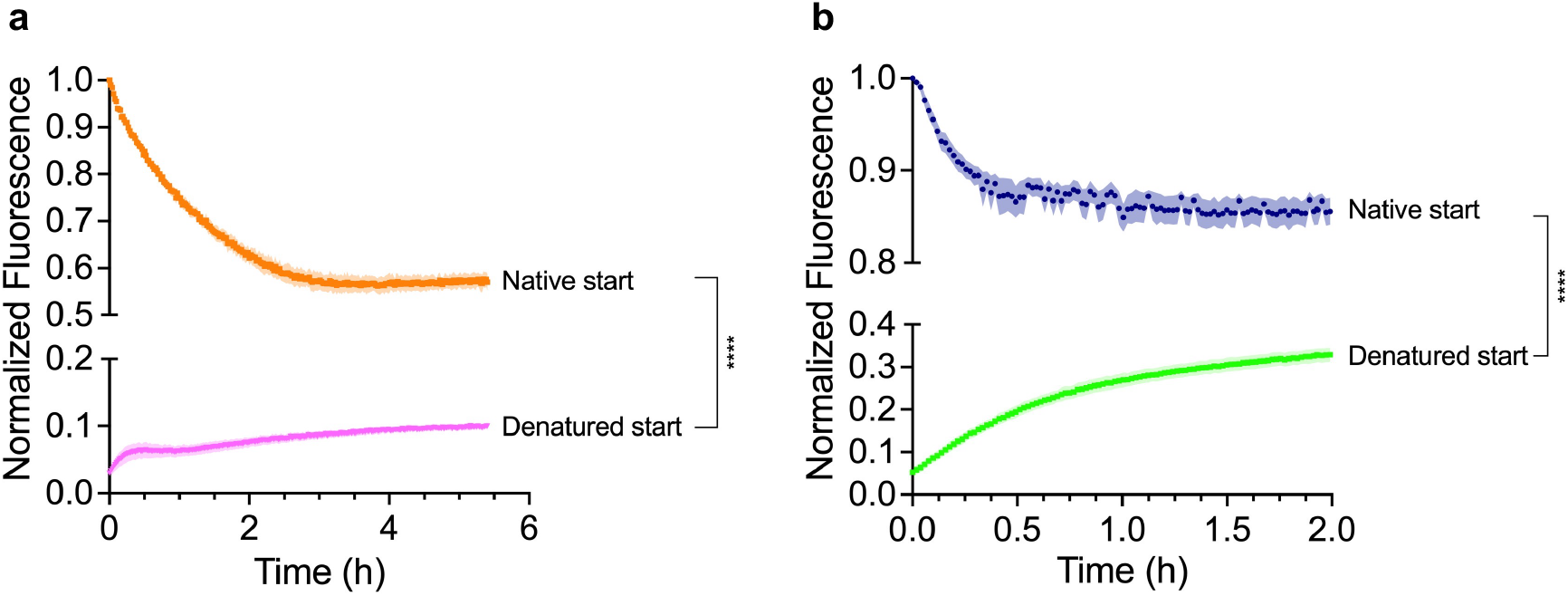
Folding after emergence out of ClpX is affected by the conformational state of the substrate pre-translocation. (**a**) ClpX-mediated translocation of native mCherry-ssrA (*orange*; n=3) leads to a higher folding yield than translocation of pre-denatured mCherry-ssrA (*pink*; n=4). (**b**) Similar results are observed for translocation of native Kaede-ssrA (*blue*; n=5) versus pre-denatured (*green*; n=4). Shading represent SEM. Fluorescence was normalized to time 0 for native start condition or to an equivalent concentration of native protein for the denatured condition. Differences between the fluorescence intensities at the final time point of each curve were analyzed using the student’s t-test (****p<0.0001).

### Implications for protein folding in the cell

Vectorial appearance is a ubiquitous feature of the cellular environment for protein folding. Amino acid sequences have presumably evolved to support proper protein folding during vectorial appearance of the protein. Yet functional requirements also constrain amino acid sequences, which may negatively impact folding efficiency. The function of each FP is to support formation of a chromophore with specific photophysical properties, a selective pressure that may conflict with the pressure to select an amino acid sequence with optimal folding properties. Vectorial appearance alone is sufficient to significantly alter FP folding outcome, including enhancing folding yield far above the yields upon dilution of the protein from a chemical denaturant. These results demonstrate that folding during vectorial appearance can preferentially populate folding intermediates that are more likely to fold to the native energy structure, versus misfolding. Consistent with this model, vectorial appearance that led to enhanced folding – including slower rates of appearance through ClpX – also led to a greater amplitude for a faster exponential phase in the observed kinetics.

It is noteworthy that the largest folding enhancement we observed was upon C-to-N-terminal appearance of Kaede, which folds in the cytoplasm after N-to-C-terminal synthesis^49^. These results suggest that there may be other constraints on the selection of amino acid residues within the Kaede N-terminus that run counter to supporting efficient folding from N-to-C-terminus. Consistent with this model, it is known that the cellular folding yield for some proteins can be quite low (e.g., the folding yield for newly synthesized wild type CFTR has been estimated at only ∼20%^67^), which may be due in part to preferential C-to-N-terminal folding mechanisms^68,69^. Presumably, proteins with functional constraints that run counter to the selection of amino acid sequences with efficient

N-to-C-terminus folding properties will be prime targets for folding assistance from molecular chaperones and other components of the cellular environment^31,32^. Collectively, these results suggest that the amino acid sequence of a protein is under selection not only for functional criteria, including the stability of the native fold, but also to support the formation of folding intermediates that can enhance folding yield during vectorial appearance.

## Materials and Methods

### Plasmids

The pACYC-Duet-based plasmid for expressing ClpXΔN_3_ with an N-terminal His-tag and the pQE-70 plasmid for expressing ClpP with a C-terminal His-tag were generously provided by Mark Akeson (University of California Santa Cruz). The pET-24a(+)-Kaede plasmid was generously provided by Justin M. Miller (Middle Tennessee State University). mCherry and SpyCatcher expression plasmids were purchased from Addgene (plasmids #54630 and #102829, respectively). Fast cloning^70^ was used to insert SpyTag at the N-terminus of mCherry and Kaede and to insert ssrA (AANDENYALAA) at the C-terminus of mCherry and SpyCatcher. The Kaede plasmid included a variation of the C-terminal ssrA tag (ADSHQRDYALAA) previously shown to be functionally equivalent to the ssrA present in mCherry and SpyCatcher^41,54^. The C-terminal ssrA of Kaede was deleted via fast cloning by removal of the -HQRDYALAA region containing the functional -ALAA motif.

### Protein expression and purification

SpyCatcher constructs were expressed in *E. coli* BL21(DE3)pLysS. An overnight culture was diluted 1:50 into flasks containing fresh Luria Broth (LB) media plus 100 μg/mL ampicillin (Amp) and 35 μg/ml chloramphenicol (CAM). The culture was incubated at 37°C with gentle shaking until an optical density (OD_600_) of ∼0.5. Expression was induced with 0.5 mM isopropyl β-D-1-thiogalactopyranoside (IPTG), and cultures were incubated with shaking at 30°C for 4 h. Cells were harvested by centrifugation (5,180x*g*, 15 min, 4°C) and pellets were stored at -80°C. For Ni^2+^ affinity purification, cell pellets were thawed and resuspended in SpyCatcher wash buffer (50 mM Tris pH 7.5, 500 mM NaCl, 20 mM imidazole) supplemented with 1 mM ethylenediaminetetraacetic acid (EDTA), 5 mM β-mercaptoethanol (BME), and a cOmplete protease inhibitor cocktail tablet (Roche). Resuspended pellets were incubated at room temperature for 15 min with gentle stirring, followed by sonication. The lysate was clarified via centrifugation (39,200x*g*, 30 min, 4°C) and the supernatant was filtered through a 0.22 µm syringe filter (Millipore) and loaded onto a 5 mL HisTrap HP column (Cytiva) equilibrated with wash buffer. After loading and column washing, protein was eluted with SpyCatcher elution buffer (50 mM Tris pH 7.5, 500 mM NaCl, 500 mM imidazole).

Wild type ClpX is a hexamer; however, for this work, ClpXΔN_3_ was used. ClpXΔN_3_ consists of three fused ClpX monomers, each lacking the 60 aa N-terminal domain^71^, that dimerizes to form the active translocation pore^71^. ClpXΔN_3_ retains the translocation activity of wild type ClpX but is more amenable to purification and assembly at low concentrations^72^. ClpXΔN_3_ was expressed in *E. coli* BLR(DE3) as described^72^, with the following exceptions: For Ni^2+^ affinity purification ClpX wash buffer (50 mM Tris pH 8, 300 mM NaCl, 100 mM KCl, 10% v/v glycerol, 10 mM imidazole) was used and the protein was eluted with ClpX elution buffer (50 mM Tris pH 8, 300 mM NaCl, 100 mM KCl, 10% v/v glycerol, 500 mM imidazole). Pooled fractions containing ClpXΔN_3_ were further purified by size exclusion chromatography, using a column (HiLoad Superdex 200) equilibrated in PD buffer (25 mM HEPES pH 7.6, 100 mM KCl, 20 mM MgCl_2_, 10% v/v glycerol).

Expression of ClpP in *E. coli* BL21 was performed as described above for SpyCatcher, with the following differences: 100 μg/mL of Amp was added to the LB media, cells were induced at OD_600_ ∼0.6-0.8 and then incubated at 30°C for 3 hours with gentle shaking. Purification was performed as described for ClpXΔN_3_, however the SEC purification step was omitted.

Kaede and mCherry constructs were expressed in *E. coli* BL21(DE3) and Top10, respectively, as described for SpyCatcher with the following differences: For mCherry, expression was induced with 0.2% arabinose (v/v), and Kaede was grown in LB media supplemented with 35 μg/ml kanamycin (Kan) and then incubated at 18ºC for ∼24 hours post-induction. Harvested cells were resuspended in either mCherry wash buffer (50 mM Tris pH 7.5, 500 mM NaCl, 20 mM imidazole) or Kaede wash buffer (50 mM Tris pH 8.3, 400 mM NaCl, 10 mM imidazole, 30% v/v glycerol), accordingly. Cells were lysed as described above for SpyCatcher, and both FPs were purified after linkage to SpyCatcher as described below.

### SpyCatcher-Spytag linkage reaction

SpyCatcher or SpyCatcher-ssrA was added to each FP at a 2:1 FP:SpyCatcher molar ratio and incubated at room temperature for 1.5 h. Afterward, SpyCatcher-tagged FPs were purified via Ni^2+^ affinity chromatography using a HisTrap HP column. Since SpyCatcher is the only protein containing a His-tag, only fluorescent proteins (FP) covalently attached to SpyCatcher were retained on the column (**Fig. S2**). Purification proceeded as described for SpyCatcher with the following exceptions: columns were equilibrated with the wash buffers corresponding to each FP noted above. SpyCatcher-mCherry was eluted with mCherry elution buffer (50 mM Tris pH 7.4, 300 mM NaCl, 250 mM imidazole). In the case of SpyCatcher-Kaede, the column was washed with binding buffer A (50 mM Tris pH 8.3, 400 mM NaCl, 10 mM imidazole, 30% v/v glycerol) followed by a second wash with binding buffer B (50 mM Tris pH 8.3, 400 mM NaCl, 10 mM imidazole, 10% v/v glycerol) before the protein was eluted with Kaede elution buffer (50 mM Tris pH 7.6, 400 mM NaCl, 500 mM imidazole, 10% v/v glycerol).

### ClpX translocation assays

ClpX translocation reactions were prepared as follows: Kaede and mCherry were unfolded by boiling for 5 min at 100°C in a solution of 4 M GdmCl and 50 mM Tris (pH 7.5). The denaturant was then removed by buffer exchanging the unfolded sample into PD buffer via an AdvanceBio Spin desalting column (Agilent) per the manufacturer’s instructions. The denatured protein was eluted into a tube containing an equivalent volume of a ClpX reaction mixture in PD buffer composed of ClpXΔN_3_ dimer (2 μM), ATP (10 mM), and 2X ATP regeneration mixture (ARM; comprised of creatine phosphokinase (0.62 mg/mL) and phosphocreatine (32 mM). The final concentration of the reaction was: FP (0.5 μM), ATP (5 mM), ARM (1X), and ClpXΔN_3_ dimer (1 μM). Reaction volumes were incubated at 25ºC in a 96-well plate. To prevent non-specific adsorption of proteins to the plastic plate, wells were pre-treated by incubating in 5% BSA (w/v) for 30 minutes, followed by two washes with PD buffer. Changes in fluorescence were monitored in a Synergy H1 microplate reader (BioTek) using the following acquisition settings: mCherry excitation (587 nm) and emission (615 nm); Kaede excitation (508 nm) and emission (534 nm). ClpX-mediated translocation of native FPs was conducted as described above, with the following differences: instead of denaturation, the native FP was mixed with the other components of the ClpX reaction to final concentration of FP (0.5 μM), ATP (5 mM), ARM (1X), and ClpXΔN_3_ dimer (1 μM).

The ClpXP degradation reaction was prepared as described above, with the addition of ClpP_14_ at a final concentration of 3 μM and incubation at 25ºC. Degradation was monitored via loss of intensity of the Coomassie-stained SDS-PAGE gel band corresponding to the substrate protein (**Fig. S5**), as quantified using ImageJ.

### Data analysis

The following equations were used for data fitting:

Single exponential:

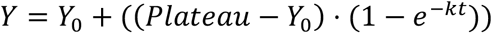

Double exponential:

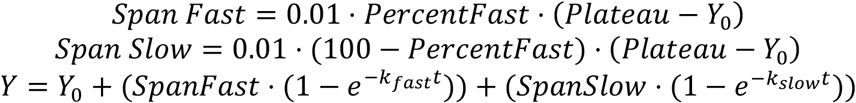

## Acknowledgments

The authors are grateful for technical advice and suggestions received from Jeff Nivala, Qing Luan and Aaron Lucius. Micayla Bowman, Esther Braselmann and Qing Luan made important technical and conceptual contributions to early prototypes for this approach. We thank all members of the Clark lab for their helpful feedback on this project. We thank the Notre Dame Biophysics Instrumentation Core Facility for technical advice and assistance.

## Author contributions

Conceptualization: PLC

Methodology: IFSS, PLC

Investigation: IFSS, PLC

Visualization: IFSS, PLC

Funding acquisition: IFSS, PLC

Supervision: PLC

Writing – original draft: PLC

Writing – review & editing: IFSS, PLC

## Data and materials availability

Plasmids constructed for this study are available upon request from the corresponding author.

