## Supplementary Information for "Protein folding success depends on the direction and speed of polypeptide chain appearance"

This file includes:

Figs. S1 to S5
Table S1
Reference 73

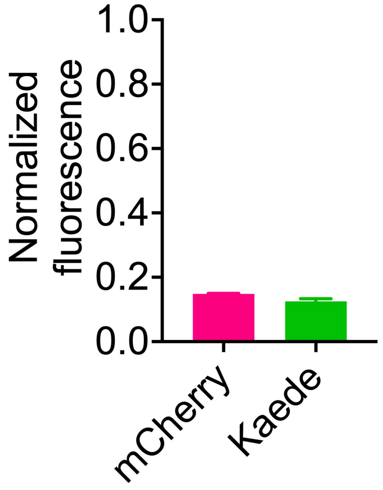


**Fig. S1.**

The fluorescent proteins mCherry and Kaede refold inefficiently upon dilution from denaturant (GdmCl). Both proteins were allowed to refold overnight at 25°C. Fluorescence emission was normalized to the signal intensity for an equivalent concentration of purified protein that was not subjected to unfolding and refolding. Columns correspond to the average of four independent experiments. Error bars represent SEM.

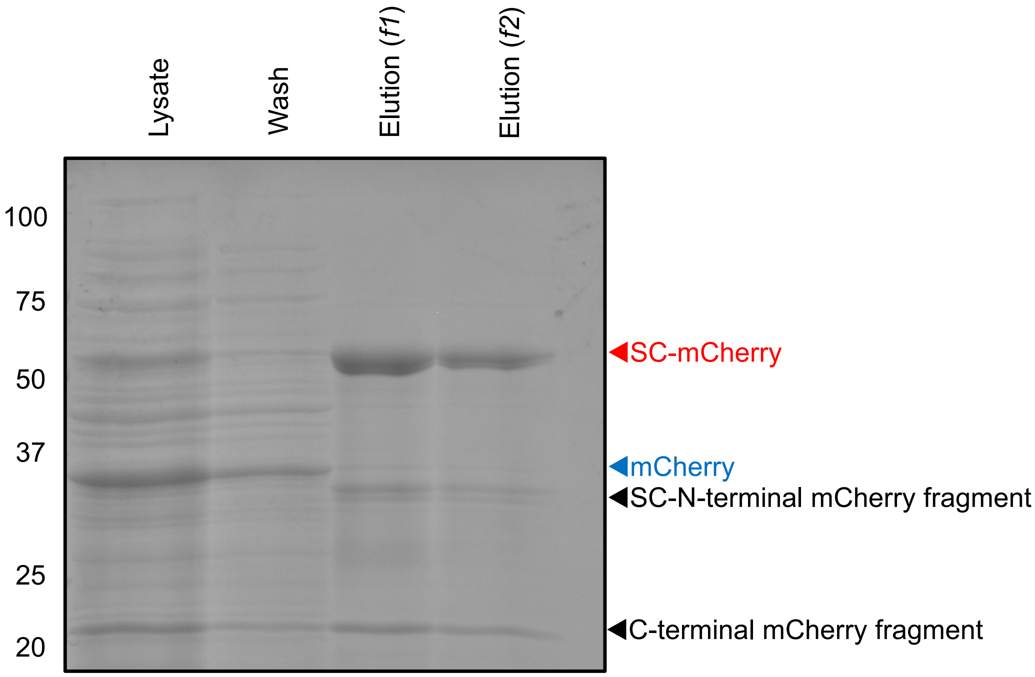


**Fig. S2.**

Purification of SpyCatcher-tagged mCherry (SC-mCherry). SDS-PAGE of fractions during $Ni^{2+}$ affinity purification. The bold band in the two elution fractions (*f1* and *f2*) corresponds to SC-linked mCherry; no band corresponding to untagged mCherry is observed. The SC-N-terminal mCherry and C-terminal mCherry fragments are the result of the cleavage of a small fraction of SC-mCherry molecules that occurs when the sample is boiled in SDS[73].

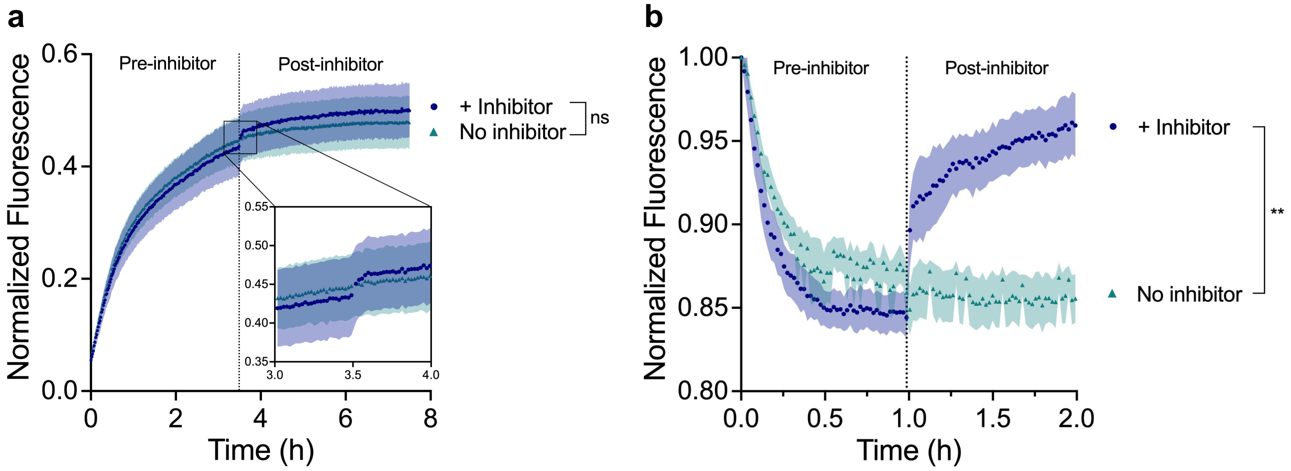


**Fig. S3.**

ClpX-mediated multiple-turnover refolding leads to a plateau in fluorescence emission intensity corresponding to a steady state between FP refolding and ClpX-mediated FP unfolding. (**a**) ClpX-mediated refolding reactions of pre-unfolded Kaede-ssrA. After 3.5 h (*dotted line*), a 60-fold excess (30 μM) of a competing ssrA-tagged substrate was added (+ Inhibitor). As expected for the addition of a competitive inhibitor of ClpX-mediated unfolding of Kaede-ssrA, this addition led to an increase in Kaede-ssrA fluorescence emission. (**b**) The fluorescence emission plateau for multi-turnover translocation of native Kaede-ssrA was also sensitive to addition of the ClpX competitive inhibitor, at 1 h (*dotted line*). In all experiments shown here, fluorescence emission intensity was normalized to the signal for an equivalent concentration of native (never unfolded) Kaede-ssrA. Data points correspond to the mean of three or five independent experiments in panels (**a**) and (**b**), respectively; shading represents SEM. Statistical analysis was conducted via Student's t-test of the fluorescence intensities at the final time point (**$p<0.01$).

**a**

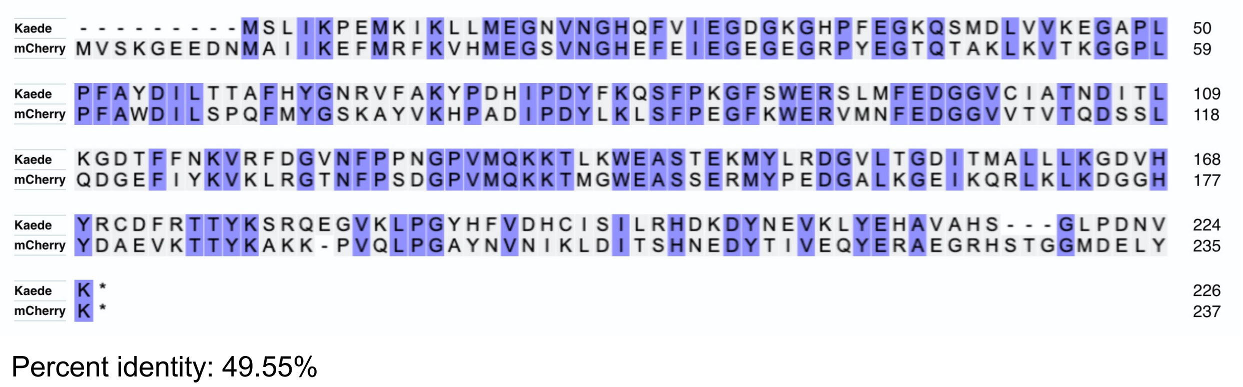


**b**

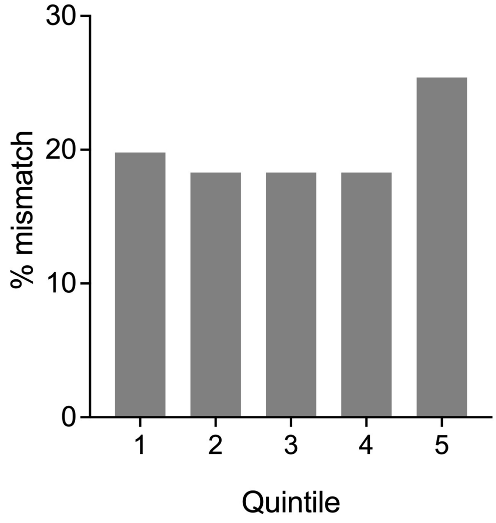


**Fig. S4.**

The sequences of Kaede and mCherry are most dissimilar at their C-termini. (**a**) Alignment of the Kaede and mCherry sequences, with identical residues indicated with *blue shading*. (**b**) Percent of mismatches occurring in each quintile of the sequence alignment, using mCherry as the reference sequence.

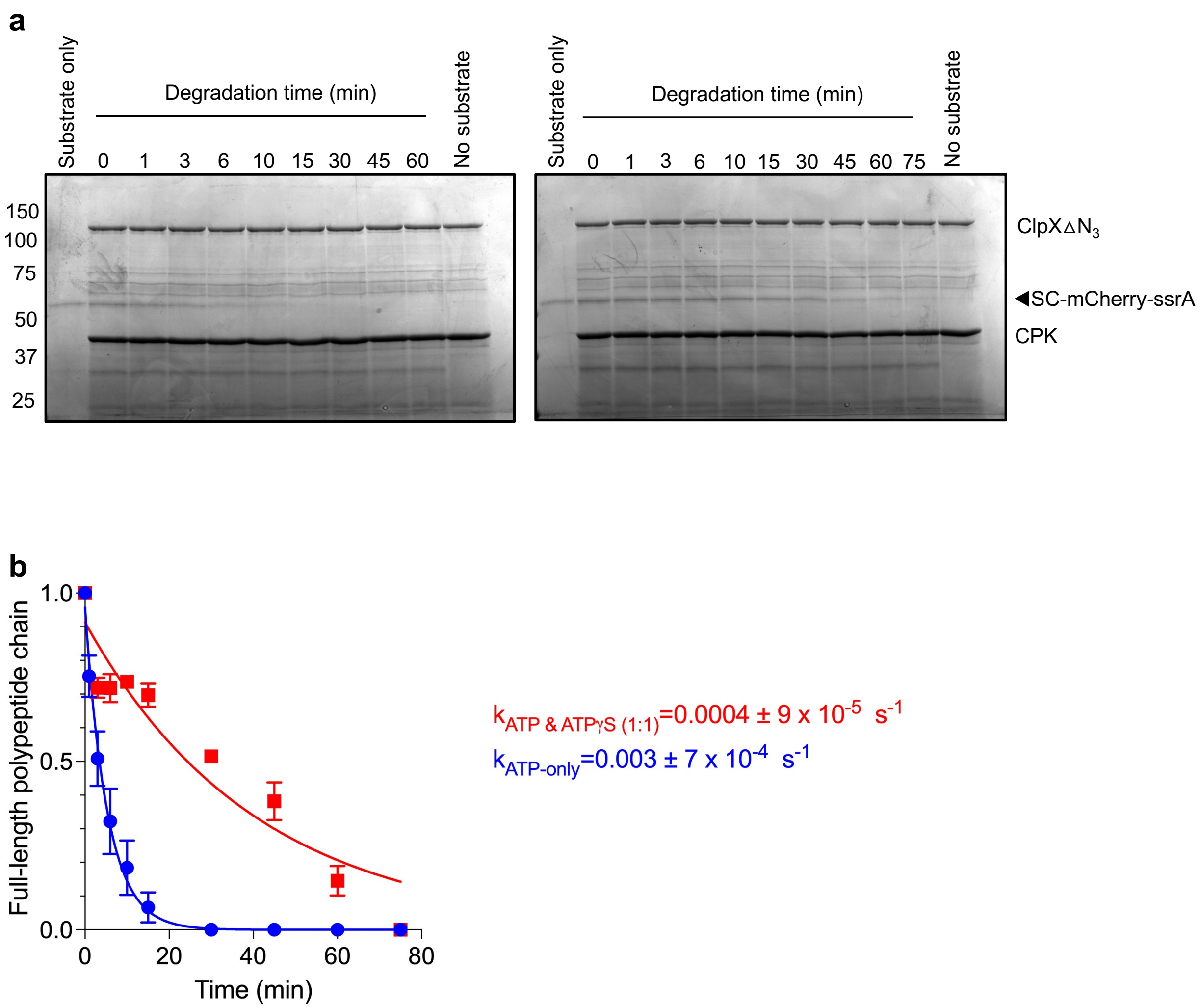


**Fig. S5.**
ClpX-mediated translocation rate is slowed down by the presence of ATPγS. (**a**) Representative SDS-PAGE gels of the ClpXP degradation of mCherry-ssrA in the presence of ATP (5 mM; *left*) or a 1:1 mixture of ATP and ATPγS (2.5 mM each; *right*). (**b**) ATPγS slows the degradation rate constant ten-fold relative to ATP alone. Each data point corresponds to the average of three independent experiments. Error bars correspond to SEM.

**Table S1.**
Kinetic parameters for mCherry and Kaede refolded under different vectorial contexts.

| Substrate | Vector | $k_{slow}$ (x $10^{-4}$ $s^{-1}$) | $k_{fast}$ (x $10^{-4}$ $s^{-1}$) | Fast phase amplitude (%) | Yield (%) |
|---|---|---|---|---|---|
| mCherry | N* | 0.7 ± 0.2 | 4.4 ± 0.9 | 70 ± 7 | 21 ± 3 |
| | N (1:1)† | 1.1 ± 0.6 | 5.7 ± 1.7 | 62 ± 13 | 22 ± 5 |
| | none (N)‡ | 0.8 ± 0.4 | 4.7 ± 3.0 | 32 ± 15 | 15 ± 1 |
| | C* | 0.9 ± 0.1 | – | – | 10 ± 1 |
| | C (1:1)† | 1.2 ± 0.2 | 13 ± 7 | 19 ± 9 | 15 ± 3 |
| | none (C)‡ | 1.3 ± 0.1 | 13 ± 8 | 21 ± 5 | 15 ± 1 |
| Kaede | N* | 1.7 ± 0.2 | – | – | 8 ± 3 |
| | N (1:1)† | 1.8 ± 0.2 | – | – | 22 ± 5 |
| | none (N)‡ | 1.4 ± 0.1 | – | – | 7 ± 1 |
| | C* | 1.4 ± 0.3 | 7.3 ± 2 | 38 ± 10 | 40 ± 5 |
| | C (1:1)† | 1.2 ± 0.9 | 3.4 ± 2 | 73 ± 27 | 51 ± 8 |
| | none (C)‡ | 1.4 ± 0.2 | – | – | 13 ± 3 |

* Vectorial appearance (N- or C- term) in the presence of ATP.
† Vectorial appearance (N- or C- term) in the presence of an equimolar amount of ATP and ATPγS.
‡ Folding in the absence of vectorial appearance. (N) and (C) correspond to the location of the ssrA tag.
Error corresponds to 95% confidence interval.
